# MEG signals reveal arm posture coding and intrinsic movement plans in parietofrontal cortex

**DOI:** 10.1101/2024.12.02.625906

**Authors:** Gunnar Blohm, Douglas O. Cheyne, J. Douglas Crawford

## Abstract

Movement planning processes must account for body posture to accurately convert sensory signals into movement plans. While movement plans can be computed relative to the world (extrinsic), intrinsic muscle commands tuned for current limb posture are ultimately needed to execute spatially accurate movements. The whole-brain topology and dynamics of this process are largely unknown. Here, we use high spatiotemporal resolution magnetoencephalography (MEG) in humans combined with a Pro-/Anti-wrist pointing task with 2 opposing forearm postures to investigate this question. First, we computed cortical source activity in 16 previously identified bilateral cortical areas (Alikhanian, et al., Frontiers in Neuroscience 2013). We then contrasted oscillatory activity related to opposing wrist postures to find posture coding and test when and where extrinsic and intrinsic motor codes occurred. We found a distinct pair of overlapping networks coding for posture (predominantly in γ band) vs. posture-specific movement plans (α and β). Some areas (e.g., pIPS) only showed extrinsic motor coding, and others (e.g., AG) only showed intrinsic coding, but the majority showed both types of codes. In the latter case, intrinsic codes appeared slightly before extrinsic codes and persisted in parallel across different cortical areas. These findings are consistent with two cortical networks for 1) direct feed-forward sensorimotor transformations to intrinsic muscle coordinates (for rapid control) and 2) computations of extrinsic spatial coordinates, possibly for use in higher-level aspects of visually-guided action, such as spatial updating and internal performance monitoring.

**Significance statement / author summary:** It is thought that the brain incorporates posture into extrinsic spatial codes to compute intrinsic (muscle-centered) motor commands, but the whole-brain temporal dynamics of this process is unknown. Here we employed human magneto-encephalography (MEG) to track this process across 16 bilateral cortical sites. We identified two, largely overlapping subnetworks for posture-dependent intrinsic codes, and extrinsic spatial coding. Surprisingly, the direct transformation from sensorimotor coordinates to intrinsic commands *preceded* the appearance of extrinsic codes, suggesting that extrinsic motor codes are derived from intrinsic codes for higher-level cognitive purposes.

## Introduction

Planning a movement toward a visual target requires transforming visual signals into appropriate muscle commands (Andersen & Buneo, 2002; Crawford et al., 2011; Shadmehr & Wise, 2005; Soechting & Flanders, 1992). This transformation needs to account for body geometry (Blohm & Crawford, 2007), effector use (Blohm et al., 2022; Scharoun et al., 2016) and arm posture (de Rugy et al., 2012). For example, the wrist muscles involved in pointing the index finger depend on hand posture: rightward pointing requires wrist *extension* with the palm facing left vs. *flexion* with the palm facing right. In both, the extrinsic (spatial) reference frame for movement was the same, whereas the intrinsic (muscle) reference frame was reversed. Humans account for such postural effects (e.g. Gielen & van Bolhuis, 1998; Lacquaniti et al., 1997; Todorov & Jordan, 2002), whether the task emphasizes extrinsic or intrinsic parameters (Kodl et al., 2011). It is generally assumed that the brain transforms sensory inputs into extrinsic motor codes^1^ and then posture is incorporated to compute intrinsic commands (Shadmehr & Wise, 2005). However, the whole-brain spatiotemporal dynamics of this process have not been clearly tested; particularly the assumption of an exclusively forward extrinsic-to-intrinsic transformation.

Most research investigating extrinsic and intrinsic motor coordinates used regional electrophysiological recordings in non-human primates. Activity of neurons in primary motor cortex (M1) depends on arm posture during reaching (Aflalo & Graziano, 2006; Caminiti et al., 1990; Scott & Kalaska, 1995, 1997; Sergio & Kalaska, 2003). M1 activity shows both muscle-specific activations (intrinsic coordinates) and movement coding (extrinsic coordinates) (Kakei et al., 1999, 2001, 2003; Morrow et al., 2007), consistent with mixed coordinate frames (Ajemian et al., 2001; Yanai et al., 2008) at an intermediate step of distributed coordinate transformations (Blohm et al., 2009). This agrees with M1 population analyses suggesting that muscle-like representations in M1 are optimized for robust control instead of reflecting direct muscle activity (Russo et al., 2018), framing M1 as a distributed population code for motor control.

Human replication attempts are sparse. A recent fMRI study has used multi-voxel pattern analysis to estimate intrinsic vs extrinsic coding across 9 parietal-frontal movement-related brain regions and found significant muscle-like (intrinsic) coordinates in M1 and S1 (primary somatosensory area), while dorsal premotor (PMd) showed extrinsic coordinates and posterior parietal cortex (likely the medial intra-parietal area, mIPS) had intermediate codes (Fujiwara et al., 2017). Consistent results have been obtained with electroencephalography (EEG) (Tanaka et al., 2018), although without the spatially selectivity. These human studies support the animal work and extend it to whole brain recordings but are limited by the trade-off between spatial and temporal resolution. Overall, the whole brain topology and dynamics of this process are unclear; in particular, the codes and relative timing of different brain areas.

Recently, we used source-localized, frequency-band specific magnetoencephalography (MEG) in humans to study the sensorimotor transformations for pointing towards or away (Pro-/Anti-pointing) from a visual target (Blohm et al., 2019), including the effector (hand) specificity (Blohm et al., 2022). Here, we combined MEG recordings with *posture-dependent* Pro-/Anti-pointing. We analyzed 16 bilateral regions of interest (ROIs) to investigate posture coding and test the timing and location of extrinsic vs. intrinsic motor codes. The above literature suggests that PMd and mIPS should show posture-independent extrinsic codes whereas M1, S1, and mIPS^2^ should show posture-dependent intrinsic codes. Our results partially confirmed these predictions; we found intrinsic-only coding in STS and SMA; mixed extrinsic/intrinsic coding in M1, PMv, S1, SPOC, aIPS and POJ; and extrinsic-only coding in aSMG, pIPS, AG and FEF. However, intrinsic motor codes generally appeared *before* extrinsic movement codes, contracting established theories.

## Methods

We chose to use MEG for its high spatio-temporal resolution (Baillet, 2017; Niso et al., 2022) to get insight into where and when posture-specific motor plans are generated in the brain and which frequencies reflect this information. We asked human participants to carry out Pro- and Anti-pointing movement in the MEG in a memory-guided task and with different forearm postures. We then co-registered MEG data with individual participant anatomical MRIs to perform whole-brain source reconstruction. We specifically extracted oscillatory activity from 16 previously identified regions of interest in the sensory-motor pathway (Alikhanian et al., 2013). We then contrasted oscillatory activity across different conditions to highlight specific coding schemes, such as posture coding, extrinsic and intrinsic movement codes. Below, we detail all these steps. Parts of these Procedures have been previously described and the dataset used was previously used for different analyses (Alikhanian et al., 2013; Blohm et al., 2019, 2022).

### Participants

10 human participants (8 males, 22-45 years old) were recruited after informed consent for a single 3h recording session involving the Pro-/Anti-pointing task with different right-hand postures in the MEG and also an anatomical MRI scan (see ‘statistics’ below for a discussion of data power). Participants were screened for history of neurological dysfunction, injury or metallic implants, and all (but one with amblyopia) had normal or corrected to normal vision. All procedures were approved by the York University and Hospital for Sick Children Ethics Boards.

### Set-up

Participants were seated Upright in a MEG scanner (151-channel, axial gradiometers, 5 cm baseline, CTF MEG system, VSM Medtech, Coquitlam, Canada, at the Toronto Hospital for Sick Children). Their head was positioned in the dewar and padded for stability if needed. Their forearm was pointing forward and supported by a forearm rest to minimize unwanted arm EMG activity leading to MEG artifacts (see Figure 1C). The forearm rest was adjustable so that pointing could be performed through wrist movements to align the index finger with the intended location on the screen. They faced a 1-m distant tangential screen used for stimulus display. Stimuli were rear-projected from outside the shielded room (Vacuumschmelze Ak3b) onto a translucent screen at 60Hz (Sanyo PLC-XP51 LCD projector with Navitar model 829MCZ087 zoom lens) using Presentation (Neurobehavioural Systems, Inc., Albany, CA, USA). Presentation also generated event timing signals that were recorded by the MEG hardware through parallel port communication. Noise levels in the room were below 10 fT/√Hz above 1.0 Hz. MEG data were online low-pass filtered at 200 Hz using synthetic third-order gradiometer noise cancelation.

**Figure 1:**
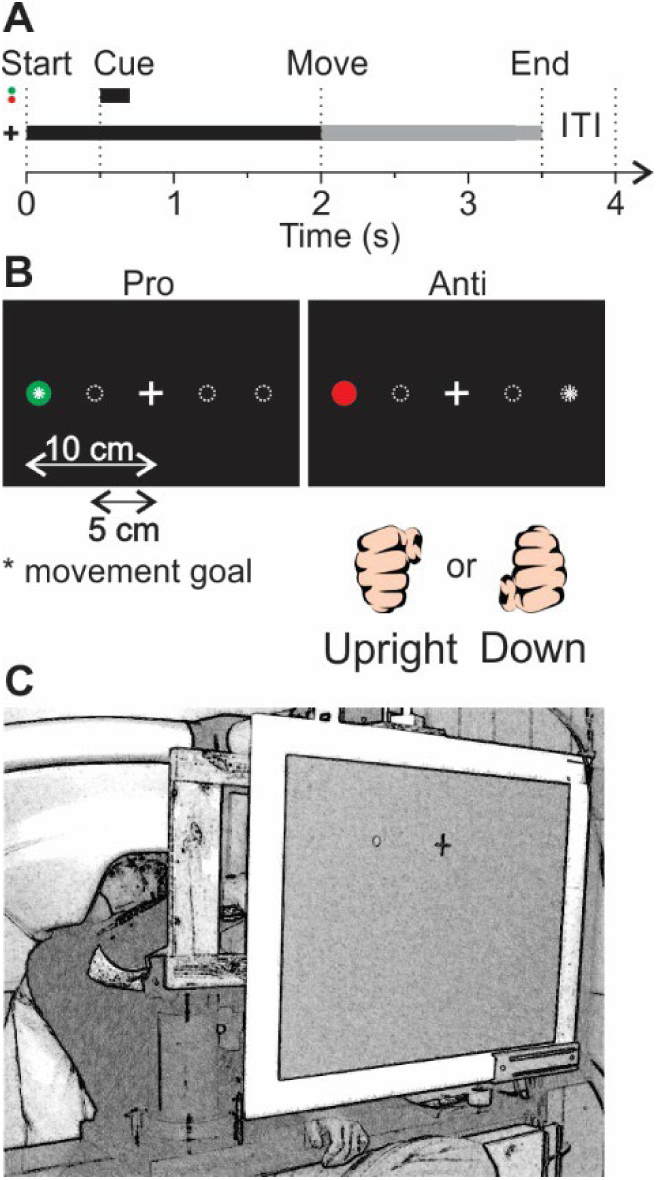
Experimental set-up. **A**. Timeline. Each trial started with a central fixation cross. 500ms later, a colored spatial cue was presented for 200ms. The cue color indicated whether a given trial was a Pro- or Anti-pointing trial (see panel B). The extinction of the cue was followed by a 1,300ms delay after which the central fixation cross was dimmed, indicating to participants to point to the goal location. Participants had 1,500ms to execute the pointing movement, followed by a 500ms inter-trial interval (ITI). **B**. Spatial arrangement on screen. Participants fixated and pointed to the central fixation cross. The spatial cues were presented at one of 4 eccentric target locations. **C**. Sketch of setup. Participants sat Upright with their head in the dewar and their forearm supported to reduce shoulder muscle activity. A wooden frame contained light barriers to help with movement detection. (modified from (Blohm et al., 2019, 2022))

To control for eye fixation, we recorded bipolar EOG; to detect wrist movement onset and direction, we simultaneously (through the MEG hardware) recorded bipolar EMG from four forearm muscles at 625Hz using Ag/AgCl solid gel Neuroline (Ambu) electrodes of type 715 12-U/C. We placed pairs of EMG electrodes over Extensor Carpi Radialis Longior (ECRL), Extensor Communis Digitorum (ECD), Extensor Carpi Ulnaris (ECU), and Supinator Longus (SL) muscles. A wooden frame mounted around the forearm contained light barrier sensors that participants would cross when pointing left or right for an independent measure of movement direction (Figure 1C).

To determine head position with respect to MEG sensors, participants were outfitted with head localization coils in the MEG. Head localization scans were performed before and after each scan (i.e. block of trials). We also obtained structural (T1-weighted, 3D-SPGR) MRI scans from a 1.5 T Signa Advantage System (GE Medical Systems, Milwaukee, WI) either before or after the MEG session. MRIs included the fiducial locations of the head coils for co-registration of MEG signals with brain coordinates (see below, Table 1). We used the individual participant T1-weighted MR data and BrainSuite software (Shattuck & Leahy, 2002) to derive the inner skull surface.

**Table 1:**
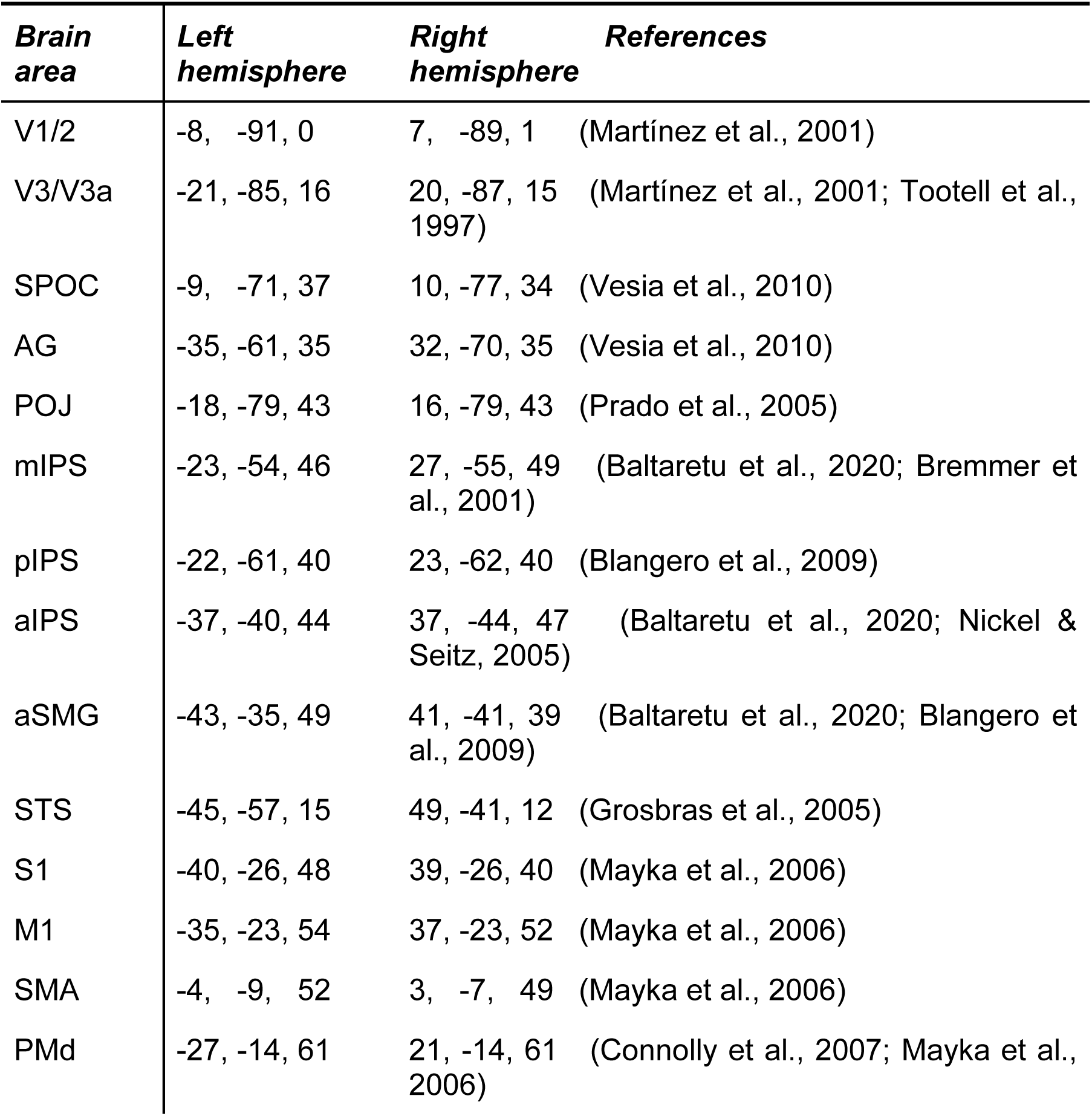
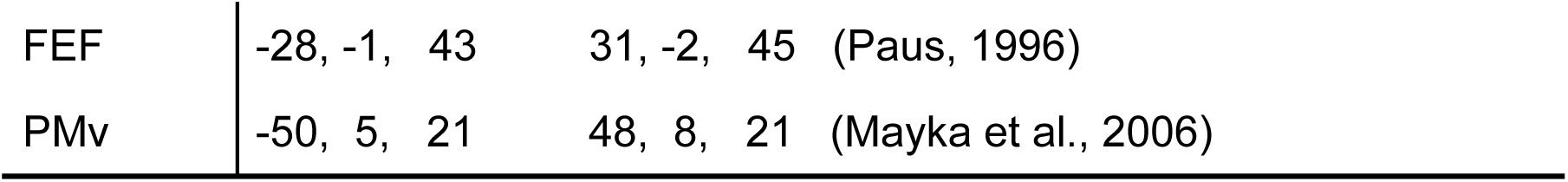
Average Talairach coordinates (mm) of functional brain sites. Activation regions of interest were identified using an adaptive clustering approach (Alikhanian et al., 2013) and cross-validated from the literature (indicated by references). We focused on sites corresponding to visual areas V1/2 and V3/3a, SPOC (superior parietal occipital cortex), AG (angular gyrus), POJ (parietal occipital junction), mIPS (medial intra-parietal sulcus), pIPS (posterior intra-parietal sulcus), aIPS (anterior intra-parietal sulcus), aSMG (anterior supramarginal gyrus), STS (superior temporal sulcus), S1 (primary somato-sensory cortex), M1 (primary motor cortex), SMA (supplementary motor area), FEF (frontal eye fields), PMd and PMv (dorsal and ventral pre-motor cortex). Note that compared to our previous studies (Alikhanian et al., 2013; Blohm et al., 2019) we updated the names of mIPS, aIPS, aSMG and pIPS to bring our nomenclature in line with the most recent literature in this area (Baltaretu et al., 2020; Blohm et al., 2022; Cappadocia et al., 2018).

| **Brain area** | **Left hemisphere** | **Right hemisphere** | **References** |
| --- | --- | --- | --- |
| V1/2 | -8, -91, 0 | 7, -89, 1 | (Martínez et al., 2001) |
| V3/V3a | -21, -85, 16 | 20, -87, 15 | (Martínez et al., 2001; Tootell et al., 1997) |
| SPOC | -9, -71, 37 | 10, -77, 34 | (Vesia et al., 2010) |
| AG | -35, -61, 35 | 32, -70, 35 | (Vesia et al., 2010) |
| POJ | -18, -79, 43 | 16, -79, 43 | (Prado et al., 2005) |
| mIPS | -23, -54, 46 | 27, -55, 49 | (Baltaretu et al., 2020; Bremmer et al., 2001) |
| pIPS | -22, -61, 40 | 23, -62, 40 | (Blangero et al., 2009) |
| aIPS | -37, -40, 44 | 37, -44, 47 | (Baltaretu et al., 2020; Nickel & Seitz, 2005) |
| aSMG | -43, -35, 49 | 41, -41, 39 | (Baltaretu et al., 2020; Blangero et al., 2009) |
| STS | -45, -57, 15 | 49, -41, 12 | (Grosbras et al., 2005) |
| S1 | -40, -26, 48 | 39, -26, 40 | (Mayka et al., 2006) |
| M1 | -35, -23, 54 | 37, -23, 52 | (Mayka et al., 2006) |
| SMA | -4, -9, 52 | 3, -7, 49 | (Mayka et al., 2006) |
| PMd | -27, -14, 61 | 21, -14, 61 | (Connolly et al., 2007; Mayka et al., 2006) |
| FEF | -28, -1, 43 | 31, -2, 45 | (Paus, 1996) |
| PMv | -50, 5, 21 | 48, 8, 21 | (Mayka et al., 2006) |

### Task

Participants were instructed to carry out a Pro-/Anti-pointing task with their right wrist while their forearm was supported in 3 different postures on separate blocks of trials: Pronation (palm facing down), Upright (thumb up, palm facing left) and Downward (thumb down, palm facing right). (Note: here we only used Upright and Downward posture data to compute extrinsic vs intrinsic motor codes; see Analysis). Each block contained 400 trials equally divided (and counterbalanced) into right and leftward cues and Pro- and Anti-conditions. The color of the spatial target cue indicated whether a given trial was a Pro-trial (point at the goal) or an Anti-trial (point to the mirror opposite) and color association (red/green) was randomized across participants. Figure 1 depicts the task details. Participants were instructed to maintain central fixation (fixation cross) throughout the trial. At the beginning of the task, participants had to point to the central fixation cross. Next, the spatial cue (Figure 1B) was presented for 200ms, followed by a 1,300ms delay (Figure 1A). The spatial cure could appear at one of four locations (left/right, 2 different eccentricities). Different eccentricities were chosen to reduce automatic pointing and encourage participants to compute an actual motor plan. We averaged data across eccentricities for the same side in the data analysis. At the end of the delay, the fixation cross was dimmed (go cue), indicating to participants to execute their Pro- or Anti-pointing movement. Participants pointed centrally during the inter-trial interval.

### Analysis

All analyses were done in Matlab (The Mathworks, Inc., Natick, MA, USA).

#### EMG and movement processing

We used EMG to precisely detect wrist movement onset. EMG was band-pass filtered (15-200Hz) and full-wave rectified. A custom algorithm then detected when EMG exceeded 3 standard deviations of baseline activity (measured before target cue onset) for each muscle. We considered the first detection time across all muscles as movement onset and this was visually inspected (and corrected when necessary: <2% of trials). We removed trials with movement direction errors (3.2% of total across participants) from further analysis. Errors were movements going into the wrong direction, i.e. opposite to the Pro-trial cue or toward the Anti-trial cue. We did not allow of later error corrections, i.e. trials were removed if initial movements were erroneous. Movement direction was confirmed by the light barriers signaling that fingers interrupted the light beam when crossing the light barrier (wooden frame in Figure 1C).

#### MEG source estimation

We used a scalar (zero-noise gain) minimum-variance beamformer algorithm (D. Cheyne et al., 2007, 2008; Hadjipapas et al., 2005; Vrba & Robinson, 2001) for MEG source reconstruction as implemented in the Brainwave Matlab toolbox (Jobst et al., 2018). This method has been shown to achieve high localization accuracy under conditions of low to moderate signal-to-noise ratio (Neugebauer et al., 2017; Sekihara et al., 2005). All following analyses were then conducted in source space.

We carried out a region-of-interest (ROI) analysis and used previously reported independently identified regions of interest (ROIs) from the same data set (Alikhanian et al., 2013; Blohm et al., 2019, 2022). Briefly, we used adaptive clustering on peak whole-brain activations in time-averaged raw, non-contrasted data to identify reliable clusters of brain activation and determined area labels that most likely corresponded to the clusters from the literature (see references in Table 1). Note that using the raw, non-contrasted data for determining ROIs was an orthogonal approach to our condition-contrasted analyses here, making this a statistically valid approach (Kilner, 2013; Kriegeskorte et al., 2009). Resulting ROIs are summarized in Table 1. ROI coordinates were transformed into individual participant MNI coordinates using standard affine transformations (linear and non-linear warping) in SPM 8 (Penny et al., 2007). Estimated individual-trial source signals at each participant-specific ROI were then computed using the above-mentioned beamformer. Next, we computed time-frequency responses (TFRs) based on these individual trial ROI data by means of standard wavelet transforms for alpha (7-15Hz), beta (15-35Hz), low gamma (35-55Hz) and high gamma (55-120Hz) bands. We also computed average whole-brain activation maps and then projected them onto a surface mesh of an average brain (PALS-B12 atlas (Van Essen, 2005)) using Caret (Van Essen et al., 2001).

#### Analysis logic

MEG data were aligned to cue onset (−500 ms to 1,500 ms around cue onset) and movement onset (−1,500 ms to 500 ms around movement onset) and extracted for further analysis. In our study design, we exploited the fact that spatial information processing in the brain is lateralized (Van Der Werf et al., 2008). This allowed us to compute spatial contrasts that emphasized variables of interest and negated irrelevant variables. This included subtracting right from left cue trials, right from left movement trials, Downward from Upright posture trials and/or right from left cortical activation (signal power in a frequency band of interest) for a given brain region. This analysis approach is a powerful tool in electrophysiology (Kuang et al., 2016) and neuroimaging (Blohm et al., 2019, 2022; Cappadocia et al., 2017; Gertz & Fiehler, 2015) Anti-reach/pointing studies. Using these contrasts then allowed us to highlight if oscillatory brain activity was correlated with posture, was dominated by sensory or motor processing and whether posture modulated motor signals, indicating intrinsic (modulation) or extrinsic (no modulation) motor coding.

Figure 2 outlines the specific logic behind motor coding and the expected difference between intrinsic and extrinsic coding in our MEG signals. Motor coding (as opposed to sensory coding) means oscillatory power changes from baseline are similar for the same movement direction regardless of conditions leading this movement. This means that if the planned movement is coded, a left-cue Pro trial should result in similar activation than a right-cue Anti-trial and vice versa (as opposed to sensory coding where left-cue Pro and Anti-trials should have the same activation, which should be different from right-cue trials). Now, if movement direction is coded irrespective of posture (extrinsic motor code), then this motor activation pattern should be identical across upward and Downward postures. Conversely, if posture is taken into account in the motor plan (intrinsic motor code), then upward and Downward postures should result in opposite (or at least different) activation patterns for upward and Downward postures. Thus, contrasting motor codes for Upright vs Downward postures should allow us to distinguish between intrinsic and extrinsic coding (see Discussion section for limitations).

**Figure 2:**
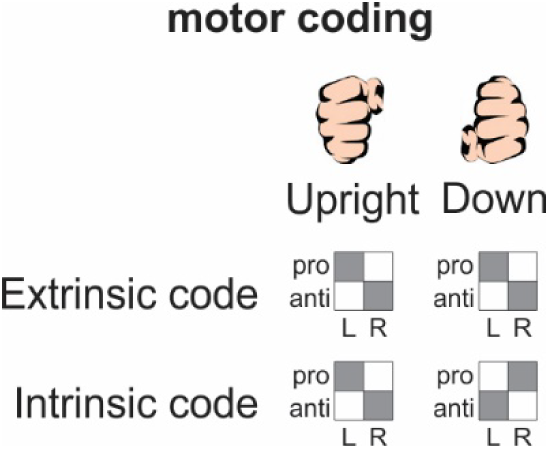
Analysis logic. Expected motor code activation patterns (frequency power) for Upright vs Downward posture and for extrinsic vs intrinsic movement plans. Patterns across Pro-/Anti-conditions and left and right cue locations reflecting motor coding should contain similar activations for the same movements, thus resulting in similar oscillatory power changes for Pro-L_cue_ and Anti-R_cue_ vs Pro-R_cue_ and Anti-L_cue_ (power changes depicted by grey and white squares). For extrinsic plans, no difference should be observed between patterns of Upright and Downward postures; however, for intrinsic plans, the pattern should flip between Upright and Downward postures because the same intrinsic (i.e. muscle activation) plan results in opposite movements for Upright vs Downward postures.

#### Contrast computations

We first wanted to investigate which sites coded for posture. To do so, we averaged data across all target conditions (left/right cure, Pro-/Anti-trials) separately for each posture and then subtracted Downward posture data from Upright posture data:

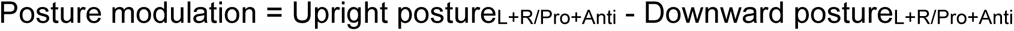

, where L/R stands for left and right cue locations (averaged across both eccentricities).

Next, we wanted to know where in the brain posture coded were integrated with motor codes. Thus, to investigate whether movement was coded in intrinsic coordinates, we first computed a motor code as follows (Blohm et al., 2019), separately for each posture:

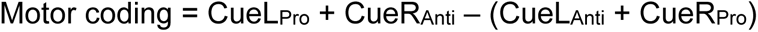

where CueL/CueR correspond to the left/right sensory stimuli presented on the screen.

Finally, to carry out goal-directed movements, the brain must specify intrinsic motor codes. To investigate whether intrinsic coding was present, we asked whether there were significant differences between motor codes in Upright vs Downward postures by computing the difference between those motor codes:

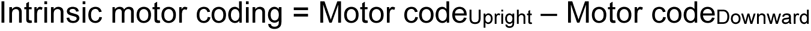

Similarly, one can compute an extrinsic motor code (independent of posture) as the sum of the motor codes across postures, i.e.

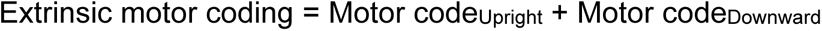

We sought to contrast the intrinsic coding results directly with extrinsic coding evaluated on the same data used for intrinsic coding. In our previous study (Blohm et al., 2019) we had computed motor coding by combining all 3 postures used (Pronation, Upright, Down). This potentially confounded intrinsic and extrinsic coding. Therefore, we only added Upright and Down postures here to evaluate extrinsic coding (instead of contrasting Upright and Down postures for intrinsic coding).

Note: we often also subtracted left from right brain site data for each ROI separately to obtain a single-site bilateral measure of posture effects and intrinsic motor codes. This is further explained in the Results section. We also checked if posture information modulated sensory coding (see Blohm et al., 2019) but did not find any significant effects.

### Statistics

We based our statistical considerations on previous neuroimaging studies (Blohm et al., 2019; Cappadocia et al., 2017; Van Der Werf et al., 2008) yielding highly robust findings with fewer participants than expected in cognitive studies. Following this literature, we designed our study with a high number of repetitions of trials to compensate for the relatively lower number of participants (Asaad & Sheth, 2024; Chaumon et al., 2021). For example, all posture modulation and intrinsic motor code computations relied on the use of ∼800 trials per participant. As in our previous studies (Blohm et al., 2019, 2022), this led to very robust results. Post-hoc power analysis (G*Power) indicated power > 0.5 (uncorrected for multiple comparisons) for alpha, beta and gamma band results in all ROIs, consistent with the standards reported in (Chaumon et al., 2021).

In order to test for statistical significance, we asked for each ROI time series of baseline-corrected frequency power whether it was significantly different from zero, i.e. significantly different from baseline. Baseline was the 500ms pre-cue period during fixation. We used temporal clustering (Maris & Oostenveld, 2007) in conjunction with 2-sided t-testing (alpha = 0.05) to evaluate significance. Temporal clustering ensured that only power changes lasting for at least 100ms were considered significant, i.e. t-test results needed to significant for at least 100ms consecutively. Significance onset timing in Figure 11 was determined through bootstrapping (N=100) across participants.

We did not perform statistical testing for whole-brain averages since these are time averages. Practically this meant that the chosen time window influenced which sites happened to appear as significant for a given time window and which ones not. This of course depended on when (and to what degree) significant activity patterns occurred during a given time window for each ROI. Thus, sometimes an ROI would show up, sometimes not, preventing a more holistic picture of the data. We therefore opted to simply display the non-significance-thresholded activations to provide a more complete picture of the data. Thus, the whole-brain projections are for visualization only.

## Results

### Overview and predictions

We analyzed data from 16 previously identified bilateral ROIs (see Table 1 and Methods), thought to play a role in planning upper limb movements. Specifically, we investigated sensory areas V1 / V2 and V3 / V3a, and S1 (primary somatosensory cortex), and higher level multisensory / sensorimotor areas SPOC (superior parietal occipital cortex), AG (angular gyrus), POJ (parietal occipital junction), pIPS (posterior intra-parietal sulcus), mIPS (medial intra-parietal sulcus), aIPS (anterior intra-parietal sulcus), aSMG (anterior supramarginal gyrus), STS (superior temporal sulcus), M1 (primary motor cortex), SMA (supplementary motor area), FEF (frontal eye fields), PMd and PMv (dorsal and ventral pre-motor cortex). We use the terms ‘sites’ to refer to the specific coordinates of these ROIs, and ‘areas’ for the surrounding regions.

Using a subset of the same dataset, we have previously shown that these brain areas are part of a dynamical network implementing the sensory-to-motor transformation for pointing (Blohm et al., 2019). For the current study we asked when and where posture information was integrated into the movement plan. In the following sections, we first analyzed posture coding by contrasting event-related activity between Upright (thumb up) and Downward (thumb Down) postures at our 16 bilateral ROIs. We then tested which of these brain areas showed posture-dependent modulations of the motor code, i.e. was the motor code different across Upright and Downward postures. The latter was then taken as being indicative of a posture-specific motor code, i.e. intrinsic motor coding. Based on the literature, we expected that a sub-network containing PMd, PMv, mIPS, SMA, and V3/V3a would show posture-independent extrinsic codes, whereas another subnetwork including M1, S1, and mIPS would show posture-dependent intrinsic coding (Fujiwara et al., 2017; Tanaka et al., 2018). Further, based on the assumption that intrinsic codes are derived from extrinsic codes, one would expect the latter codes to appear first in our dataset (Ajemian et al., 2001; Yanai et al., 2008).

### Posture modulation

To investigate where posture was coded in our network of brain areas, we first contrasted event-related data from Upright and Downward postures, averaged across all left/right Pro/Anti cue conditions (see Methods). An example result of this analysis is shown in Figure 3 for posture modulation in PMv in the high gamma band (55-120Hz) leading up to movement onset (time 0s). (Other frequency bands will be considered below.) We found that in PMv, posture significantly changed the oscillatory power in different ways for Upright (desynchronization) vs Downward (resynchronization) trials. We thus contrasted Upright and Downward hand posture to obtain the *3^rd^ row* of Figure 3. In this case, the blue data in the final combined average (lower row) indicate the relative desynchronization (i.e., relatively more activation) for the Upright hand posture. Although pointing was only done with the right hand in this study, similar modulations were observed in both left (contralateral) and right (ipsilateral) PMv (*left and middle columns* of Figure 3. Therefore, we averaged across both sides to obtain a single bilateral plot (e.g., Figure 3, 3rd column). The combination of these two comparisons (Upright – Down posture and Left + Right Hemisphere) resulted in a single measure of posture modulation for each bilateral ROI pair, as shown in Figure 3 (*lower-right panel*) for PMv. We henceforth refer to this as the *posture modulation* parameter.

**Figure 3:**
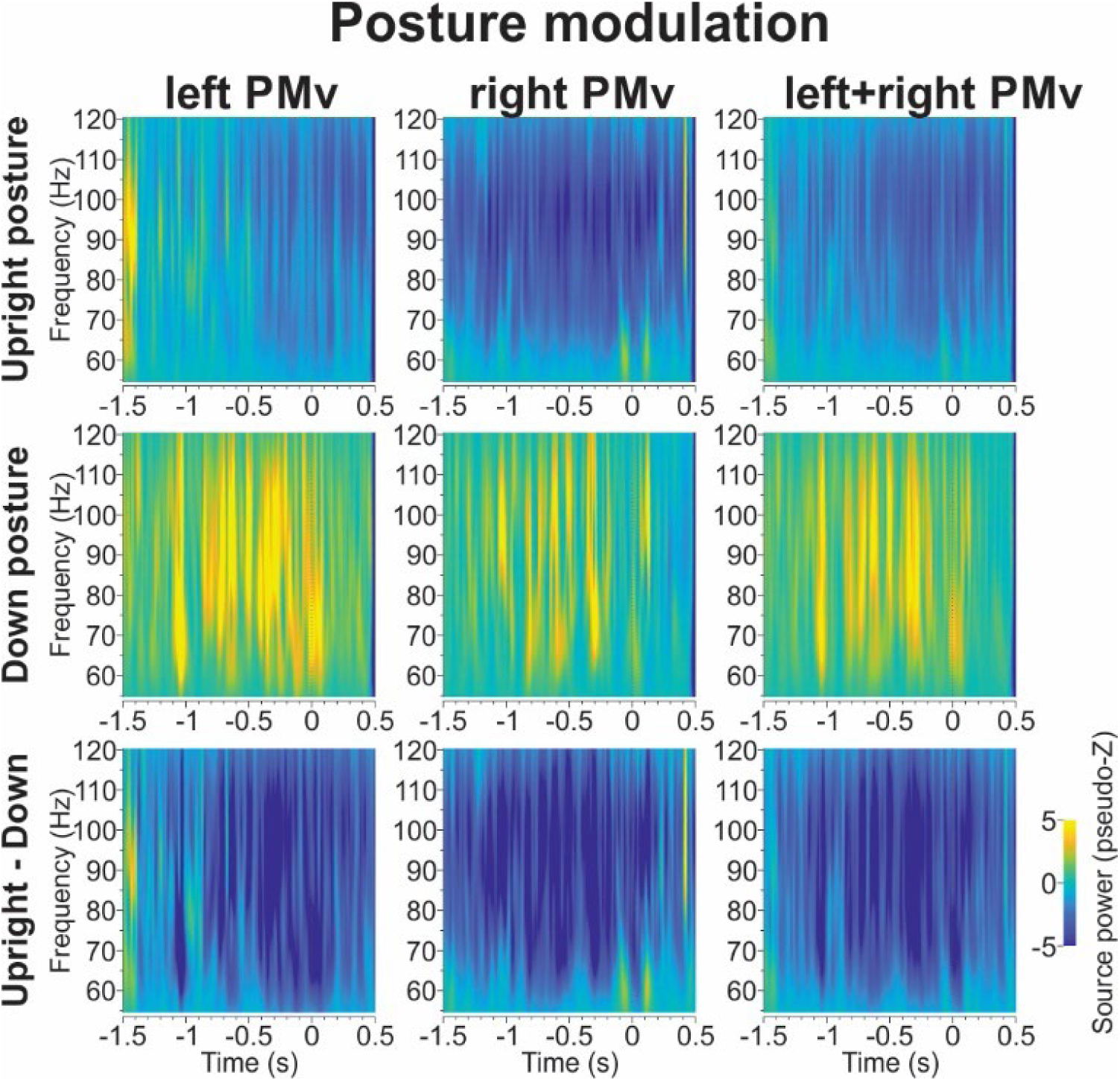
subtraction logic for example posture coding in PMv. Average movement aligned (movement onset time = 0s) change in oscillatory power for the high gamma band (55-120Hz). First and second column show results for left and right PMv, respectively. Third column shows the average PMv effect. Third column shows the Upright vs Downward posture contrast. The last panel (bottom right) represents a single bilateral index of posture modulation, i.e. Upright-Down contrast averaged across left and right PMv.

Except for SPOC, all of our ROIs showed similar bilateral modulations, so we computed posture modulation in the same way as PMv (Figure 3) for those areas. In the case of SPOC, posture modulation was opposite for the left and right hemisphere, so we subtracted right - left side data to obtain a similar measure of posture modulation.

To investigate posture coding across all ROIs, we computed posture modulation for each frequency band (alpha, beta, low gamma, high gamma) as well as for both cue-aligned and movement-aligned data. Figure 4 shows the significant posture modulations averaged across both pairs of the bilateral ROI, as explained above. We found significant posture modulation in the alpha band (SPOC), beta band (SMA, STS, FEF), low gamma band (STS, AG, PMv, FEF) and high gamma band (S1, PMv, FEF). Interestingly, all areas but SPOC (the outlier for bilateral activation) also showed Upright posture-related desynchronization. In addition, it seems that source power was consistent across much of the trial (e.g. always negative – desynchronization), even if that trend did not always reach significance. Overall, this analysis shows that a sub-set of bilateral parietal-frontal ROIs displayed significant posture modulation across different frequency bands during the delay period of the task.

**Figure 4.**
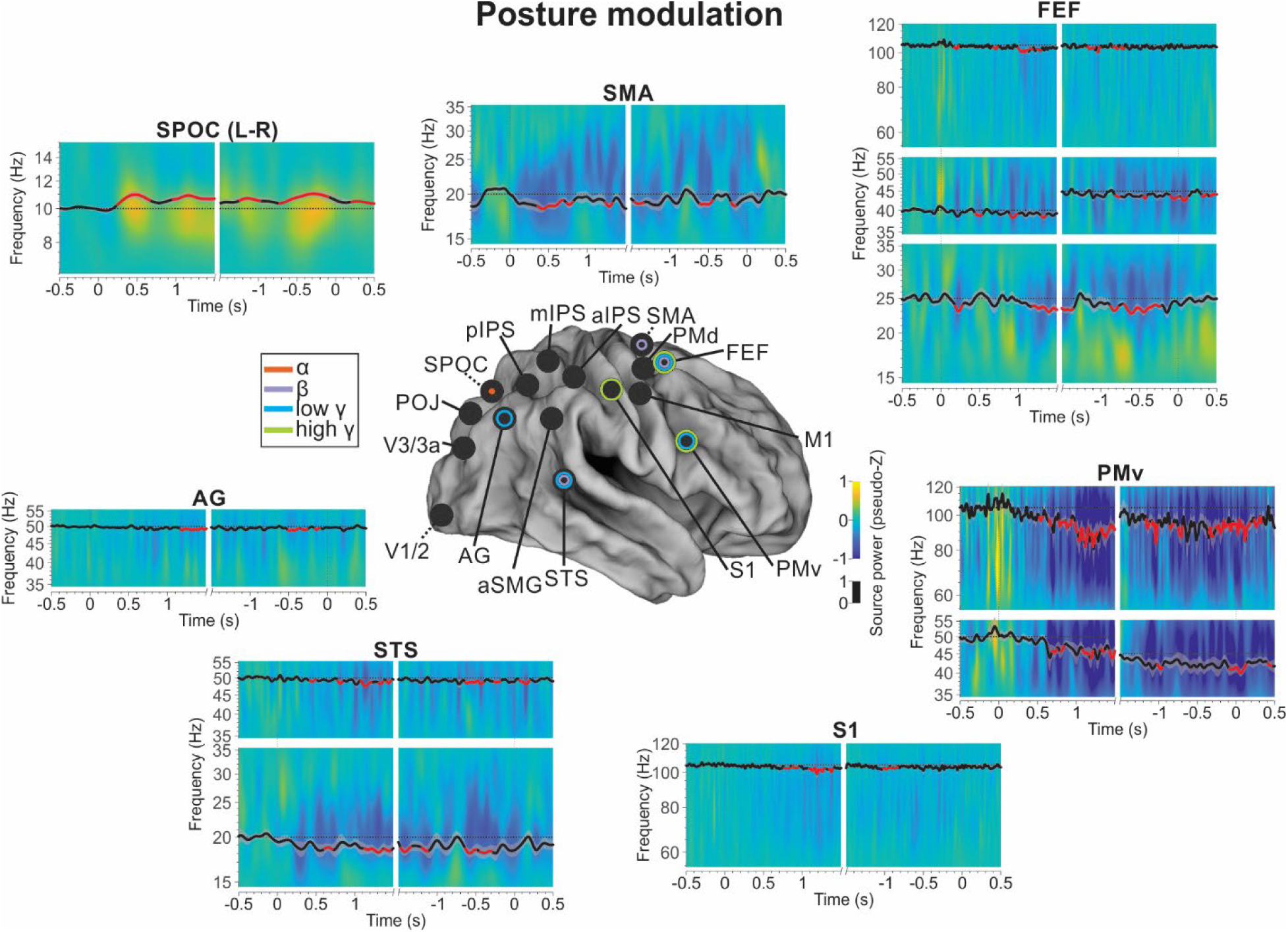
posture modulation analysis across all frequencies. Single area time-frequency response plots are shown (same as in Figure 3) with single-frequency time course overlaid. The single-frequency time course is aligned on the frequency represented (y-axis alignment, dotted horizontal line = 0) and scaled according to the black source power scaling bar (center). Black and grey traces show mean and 95% CI, and red indicates significant difference from zero. For compactness, only data from sites with significant results are shown. The center brain color codes significant sites as a function of the frequency band in which significance is found (see color key).

### Extrinsic coding results

Extrinsic movement coding means that the motor code is specified in terms of spatial movement in the environment and is therefore independent of posture. As a result, we expect to find the same motor coding pattern regardless of posture in areas showing extrinsic motor coding. The prediction is thus that the spatial tuning in the Pro / Anti task should be posture-*independent* (Figure 2). Following the same logic and conventions Figure 5 provides an example of extrinsic coding is illustrated for the left aIPS. Note that in these data, a change in hand posture from upright (A) to down (B) fails to reverse the activation pattern (in contrast to the intrinsic codes to be shown in the next section).

**Figure 5:**
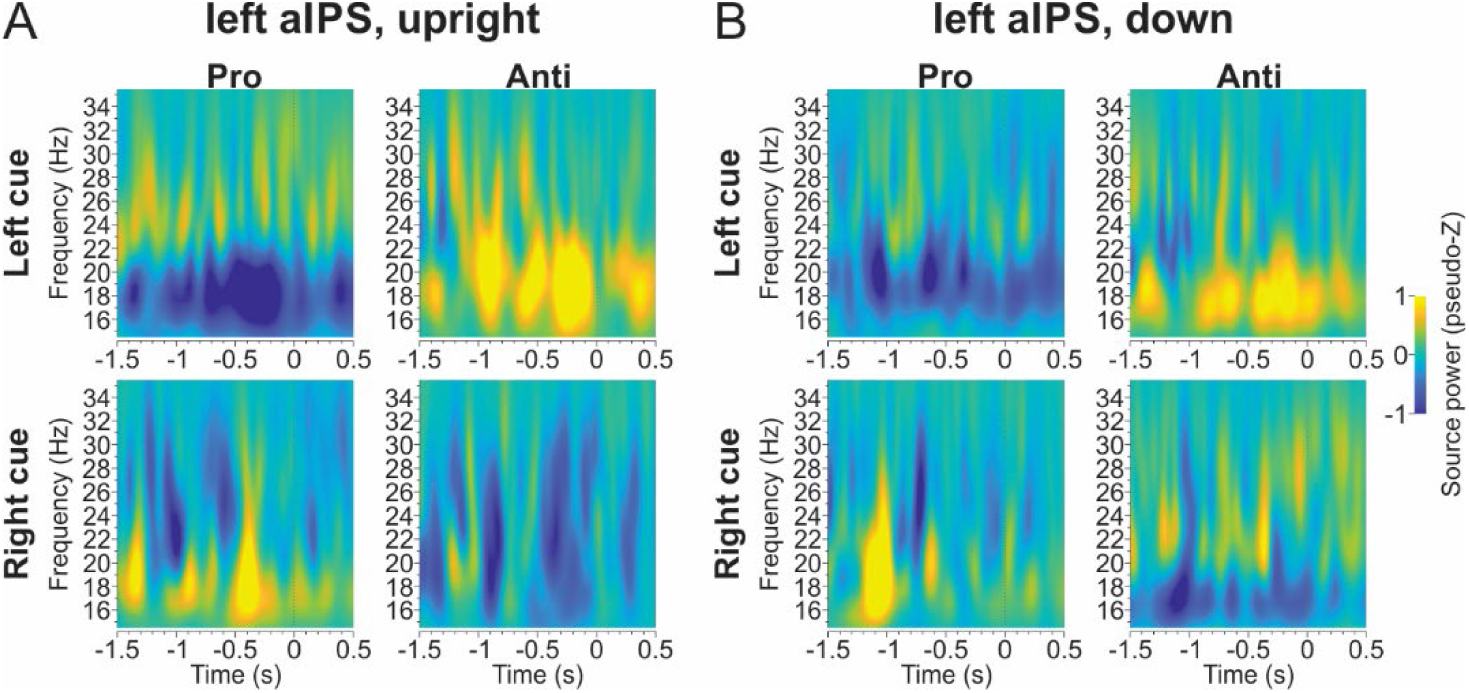
Posture-independent motor coding. Left hemisphere aIPS band oscillatory source power. Panels show Upright (A) and Downward (B) posture data. For each posture, we show the motor coding pattern, i.e. similar activity for Pro left cue and Anti right cue, which is different for Pro right cue and Anti left cue trials. This pattern of de-/re-synchronization is consistent between Upright and Downward postures, indicating extrinsic motor coding.

Figure 6 illustrates the addition/subtraction logic we used to derive a single bilateral brain area index for extrinsic motor coding. Since we expect extrinsic codes to be posture-independent, we added time-frequency responses across both hand postures, effectively averaging out any differences between postures. This can be seen in Figure 6 as the addition of the top two rows, resulting in the bottom row. Taking into account the brain’s lateralized spatial coding (Van Der Werf et al., 2008), we also subtracted right from left hemisphere data to obtain a single bilateral area-specific motor code. This can be seen in Figure 6 as the subtraction of the middle column from the left row, resulting in the right row. The lower-right panel of Figure 6 shows the combination of both of these manipulations: the extrinsic power spectrum for bilateral aIPS.

**Figure 6:**
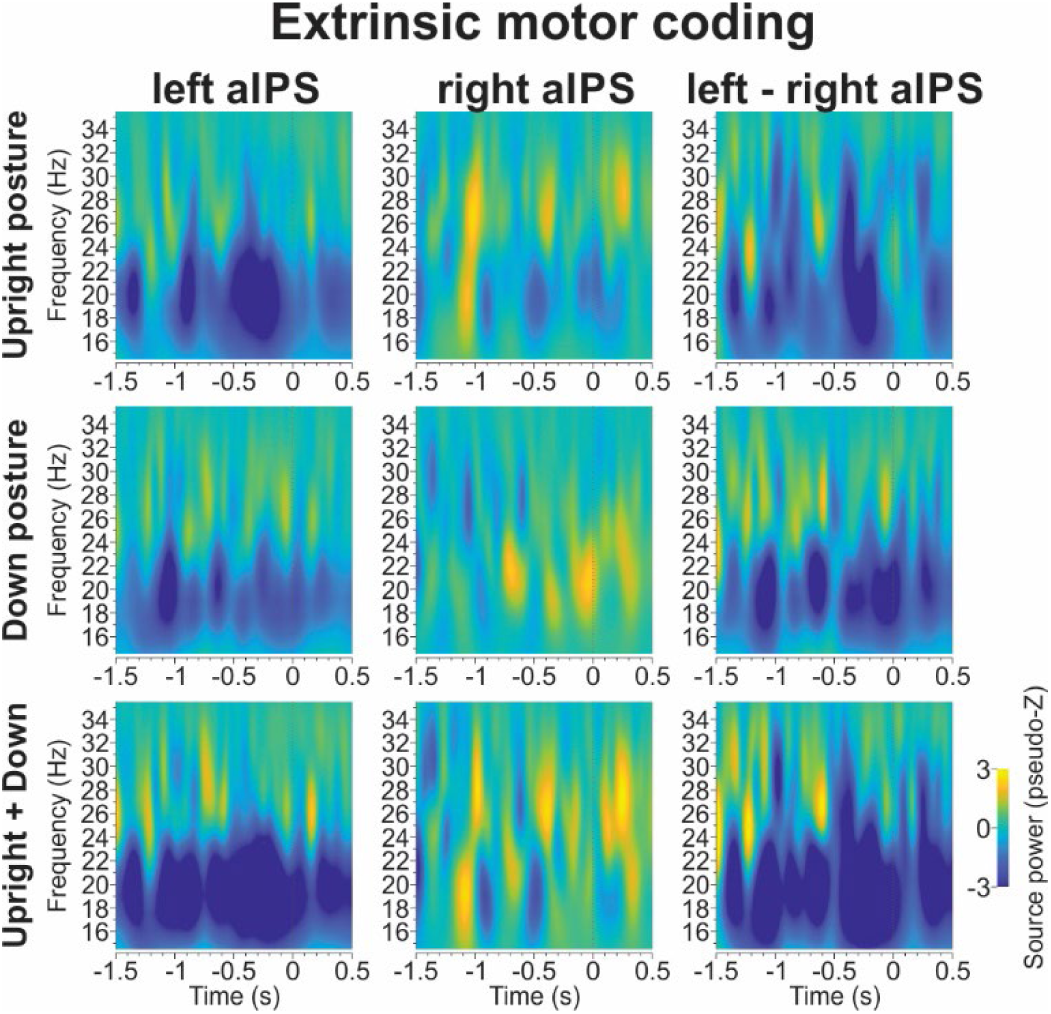
Subtraction logic for extrinsic motor coding in example area aIPS. Movement-aligned time-frequency response for bilateral aIPS. Same conventions as in Figure 3. Note that the bottom right panel represents the additive pattern of Upright and Downward postures and the difference between left and right hemispheres.

Using these conventions, we can now summarize the extrinsic codes across all of our 16 bilateral brain areas and all frequency bands (Figure 7). Interestingly, while there were some significant alpha (pIPS, POJ) and beta band (pIPS, aIPS, SPOC, AG) extrinsic codes, significant extrinsic codes were also found in low gamma (pIPS, M1, S1, AG) and high gamma bands (pIPS, aIPS, FEF, M1, PMv, S1, aSMG). The earliest occurrences of extrinsic coding happened in FEF (170ms after cue onset), followed by POJ, S1 and M1 (all ∼325ms). Few areas showed sustained extrinsic motor coding until/past movement onset, such as aSMG, S1, M1 and aIPS. Overall, we found a sub-network of extrinsic motor coding areas that includes the low and high gamma bands.

**Figure 7:**
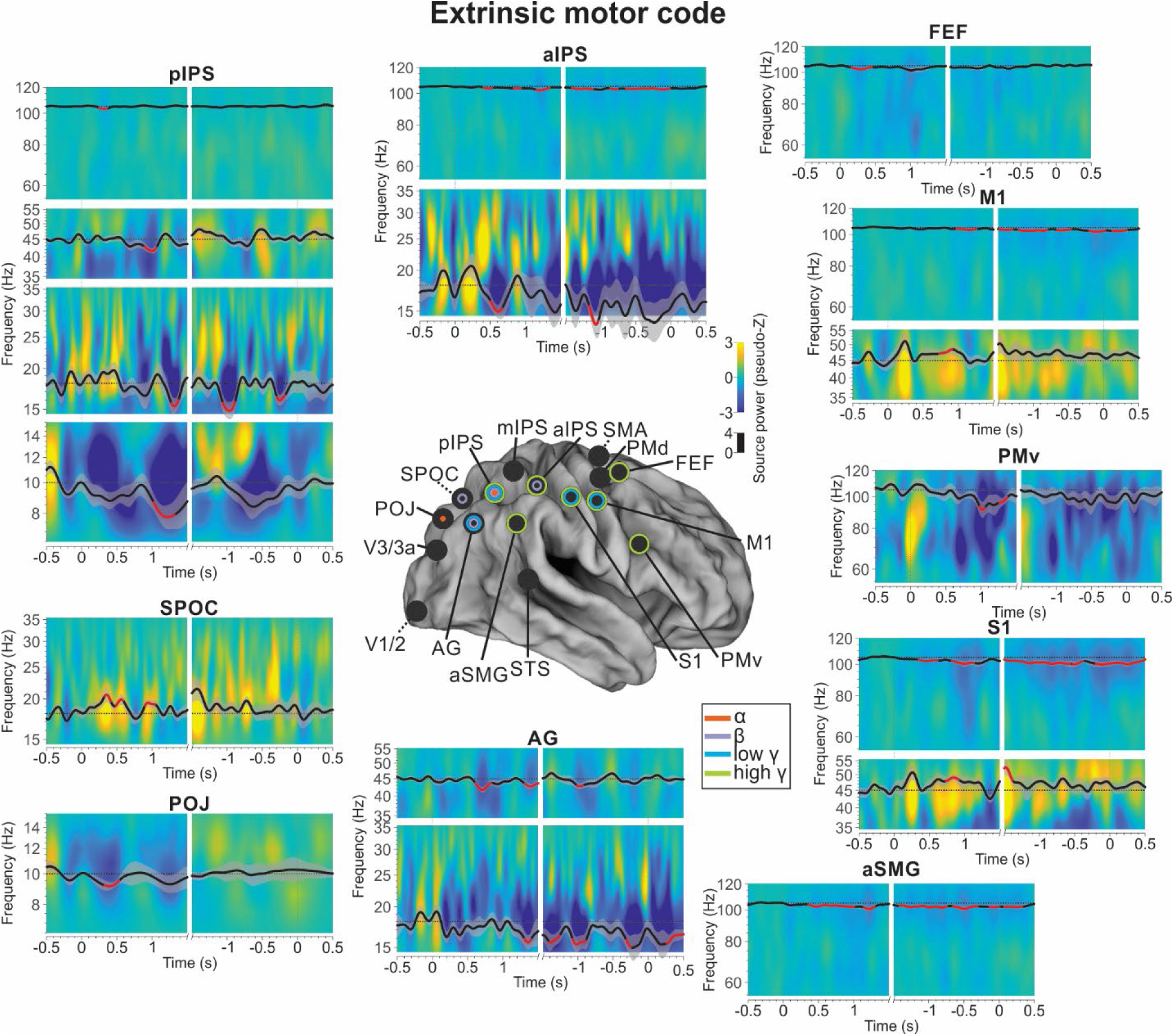
extrinsic motor coding results. Same plotting conventions as in Figure 4.

### Intrinsic motor coding

Extraction of intrinsic motor codes from extrinsic codes requires posture-specific motor computations. For example, in our right-hand task, rightward extrinsic motion requires opposite intrinsic plans from opposite hand postures: wrist extension at the Upward posture versus flexion at the Downward posture. (Conversely, the same intrinsic motion (say extension) could result in opposite extrinsic motion: rightward at the Upward posture and leftward at the Downward posture.) We used this logic to investigate intrinsic motor coding in the brain, arranging the data such that spatial tuning in the Pro / Anti should be posture-dependent (Figure 2).

We begin with an example ROI pair (M1) where the literature would clearly predict a posture-dependent, intrinsic coding scheme (Figure 8) following the logic in Figure 2. Here, we compare changes in oscillatory power separately for Pro-/Anti-trials for left cue (*upper row*) versus right cue (*lower row*) trials, and at the two forearm postures: Upright (Figure 8A) and Downward (Figure 8B). For the Upright posture (Figure 8A), right/Pro and left/Anti show desynchronization (blue, associated with activation) whereas left/Pro and right/Anti trials lead to a relative re-synchronization (yellow, associated with inhibition). This appears to suggest a code for *rightward* hand motion at this posture, but it is not yet clear whether this is coded in extrinsic or intrinsic codes.

**Figure 8.**
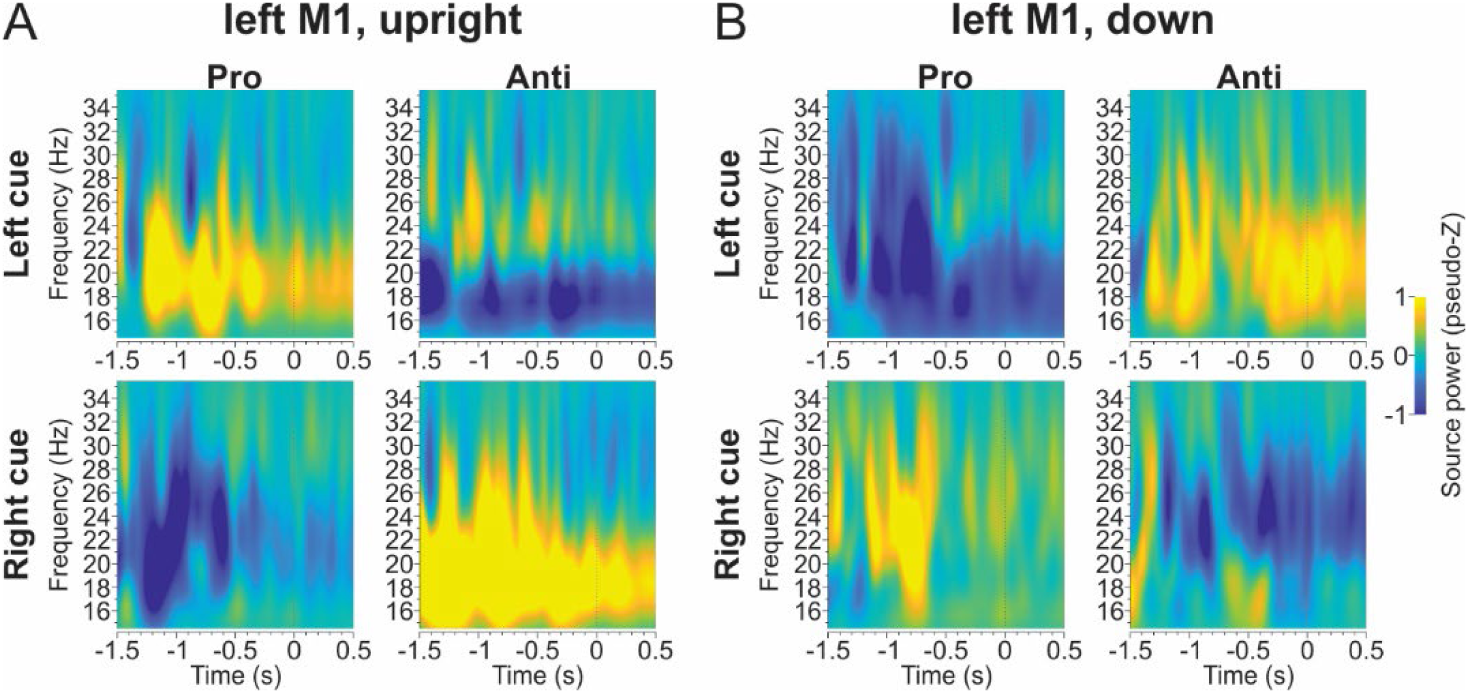
Motor coding changes with posture. Left hemisphere M1 beta band oscillatory source power. Different panels show Upright (A) and Downward (B) posture data. For each posture, we show the motor coding pattern, i.e. similar activity for Pro left cue and Anti right cue, which is different for Pro right cue and Anti left cue trials. Critically, the pattern of de-/re-synchronization is flipped between Upright and Downward postures, indicating intrinsic motor coding.

If M1 employs an extrinsic code, one should see the same pattern as Fig. 8A at the Downward posture (Figure 8B). However, the exact opposite pattern emerges: desynchronization for left/Pro and right/Anti and re-synchronization for left/Anti and right/ Pro trials. In other words, left M1 now appears to code *leftward* extrinsic hand motion. The simpler explanation is that this area codes the muscles for wrist flexion at both postures. In other words (as expected) left M1 employs an intrinsic motor code.

Figure 9 illustrates the series of subtractions that we used to derive a single measure of intrinsic (posture-dependent) motor coding for each bilateral ROI pair, again using M1 as the example. The *left column* shows how we collapsed unilateral data (Left M1 from Figure 8) from both postures into a single measure. The first (*upper-left*) panel collapses the four panels from left M1/Upright posture using the following formula: (Pro/Leftward + Anti/rightward) – (Pro/Rightward + Anti/Leftward) (see Methods). The resulting (yellow) data indicates relative desynchronization (activation) for rightward hand extension. The 2^nd^ panel (*left column, middle row*) shows the same procedure for the Downward posture, now indicating the opposite directional tuning as described above, i.e. desynchronization for leftward hand extension. Finally, the subtraction between these two (*first column, bottom row*) provides an overall measure of intrinsic coding, where extrinsic coding should cancel out. The yellow indicates relative desynchronization (activation) for extensor movements (and conversely, resynchronization / deactivation for flexor movements).

**Figure 9:**
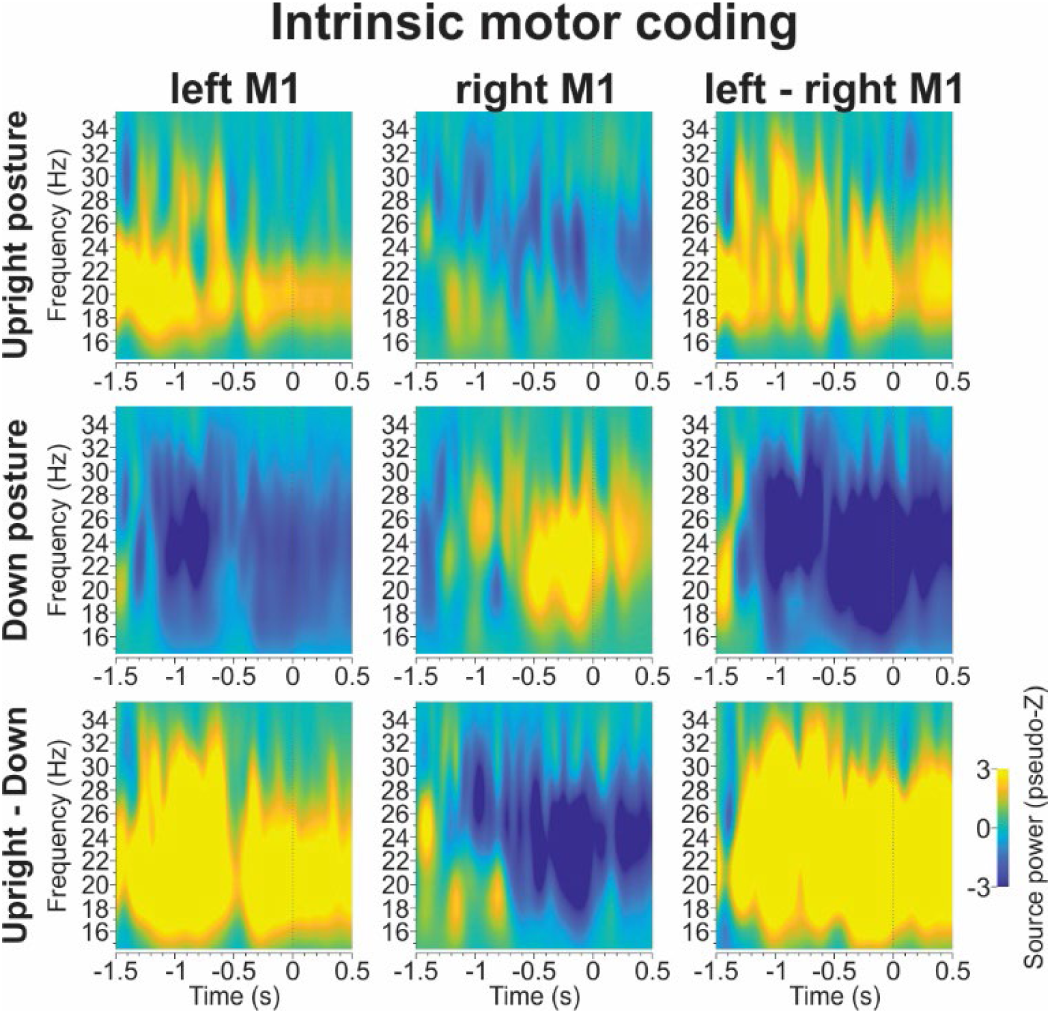
Subtraction logic for intrinsic motor coding in example area M1. Movement-aligned time-frequency response for bilateral M1. Same conventions as in Figure 3. Note that the bottom right panel represents the double-difference between Upright and Downward postures and between left and right hemispheres.

Both sides of M1 (and all other bilateral ROIs tested) showed spatially tuned oscillations for right hand motion (Blohm et al. 2019; 2022). Thus, were also able to perform an analysis of posture-dependent tuning in the ipsilateral ROIs. The *2^nd^ column* in Figure 9 repeats our subtractions from the *first column* on Right M1, resulting in the opposite frequency modulations. Finally, we subtracted the *2^nd^ column* data from the *1^st^ column* to obtain the *3^rd^ column*; the bottom right double-contrast provides a single-area summary for intrinsic motor coding, which can be interpreted as contra-lateral M1 activation (and ipsilateral M1 suppression) during wrist extension in the right hand. This result is reassuring (and not surprising) but we can now use the same test to investigate intrinsic coding in all our bilateral ROI pairs, where the coding is less obvious.

Figure 10 shows significant intrinsic motor coding across all ROIs. While posture coding in Figure 4 and extrinsic motor coding in Figure 7 were present in all frequency bands we tested, intrinsic motor coding was only found in alpha (SPOC, POJ, STS, PMv, aIPS) and beta (SPOC, SMA, M1, S1) bands. This sub-network of intrinsic motor coding areas was different but overlapped with the sub-network of posture coding. Interestingly, some areas (POJ, aIPS, STS, PMv) coded for intrinsic motor plans very early after the presentation of the cue, i.e. 186ms after cue onset for POJ, followed by S1 (235ms), PMv (290ms) and STS (315ms). Alpha band intrinsic motor coding also seemed to systematically precede beta band intrinsic motor coding. It was also interesting to see that several areas showed sustained intrinsic motor coding until/past movement onset, such as SMA, M1, S1, SPOC and STS. Overall, there was strong evidence for intrinsic motor coding in alpha and beta bands across a surprisingly distributed sub-network of parietal-frontal areas.

**Figure 10:**
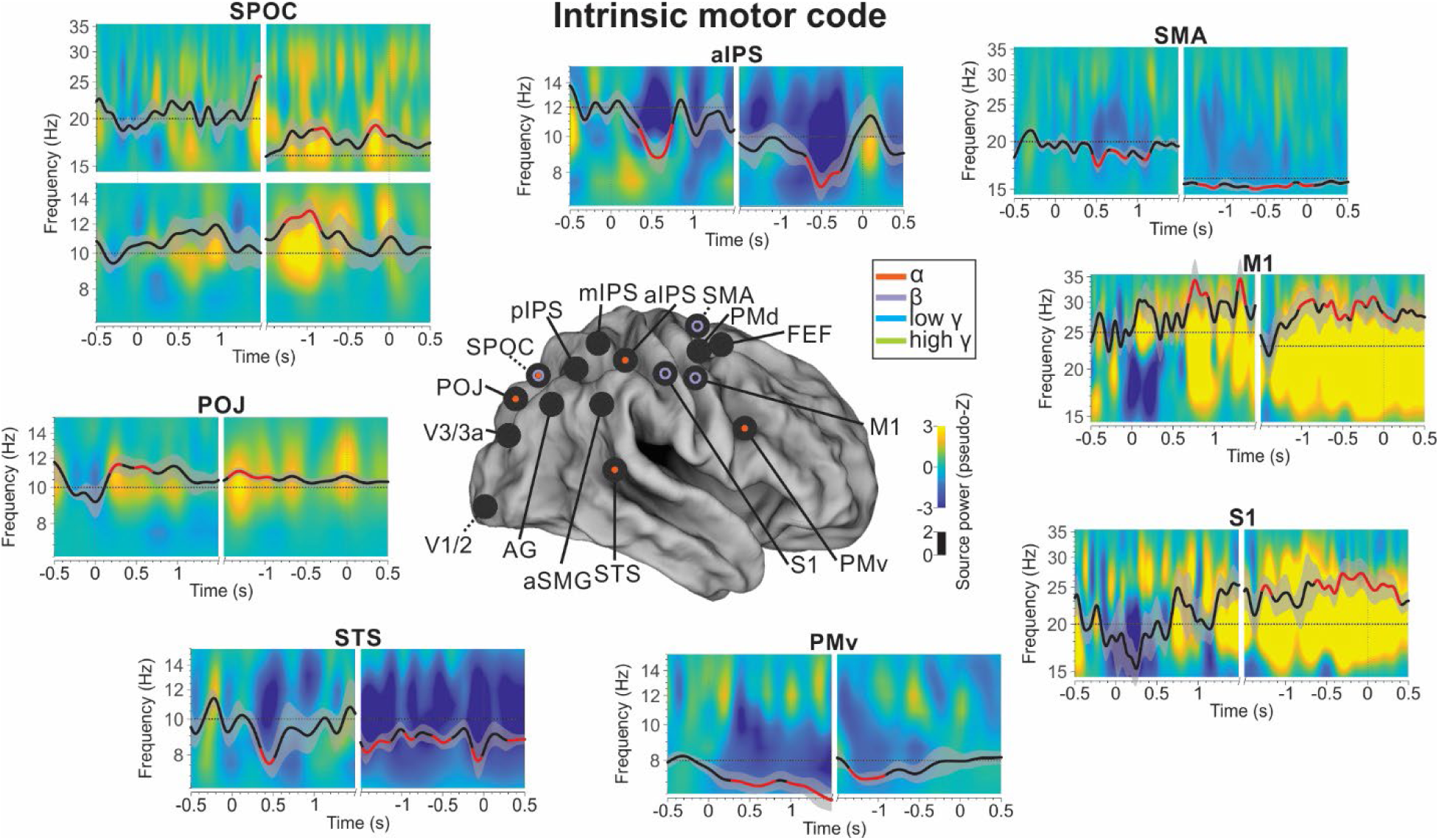
intrinsic motor coding results. Same conventions as in Figure 4.

### Summary: timing and comparison between extrinsic and intrinsic coding

Figure 11 summarizes our findings and analyzes the temporal sequences of significant posture and movement codes (both extrinsic and intrinsic). Panel A summarizes how different but overlapping sub-networks are involved in coding for posture (green), intrinsic motor plans (yellow) and extrinsic motor plans (magenta). Visual areas V1-V3 did not show significant modulations in any of these dimensions (presumably because of the motor nature of the task) with most codes appearing in parietofrontal cortex. While a few areas (SPOC, S1, PMv) show significance for all 3 codes (posture, intrinsic, extrinsic), most areas seem to be more specialized. For example, POJ, aIPS and M1 show both intrinsic and extrinsic coding, but are not significantly posture modulated. AG and FEF show posture and extrinsic motor coding, but no significant intrinsic code. SMA and STS show posture and intrinsic coding, but no extrinsic code. Finally, pIPS and aSMG show only extrinsic coding. As one might expect, posture coding was often associated with posture-dependent intrinsic codes (SPOC, STS, S1, SMA, PMv) but posture / extrinsic coding associations were just as common (SPOC, SMA, S1, FEF, PMv). Overall, parietofrontal cortex showed every possible combination between posture, intrinsic and extrinsic motor coding.

**Figure 11:**
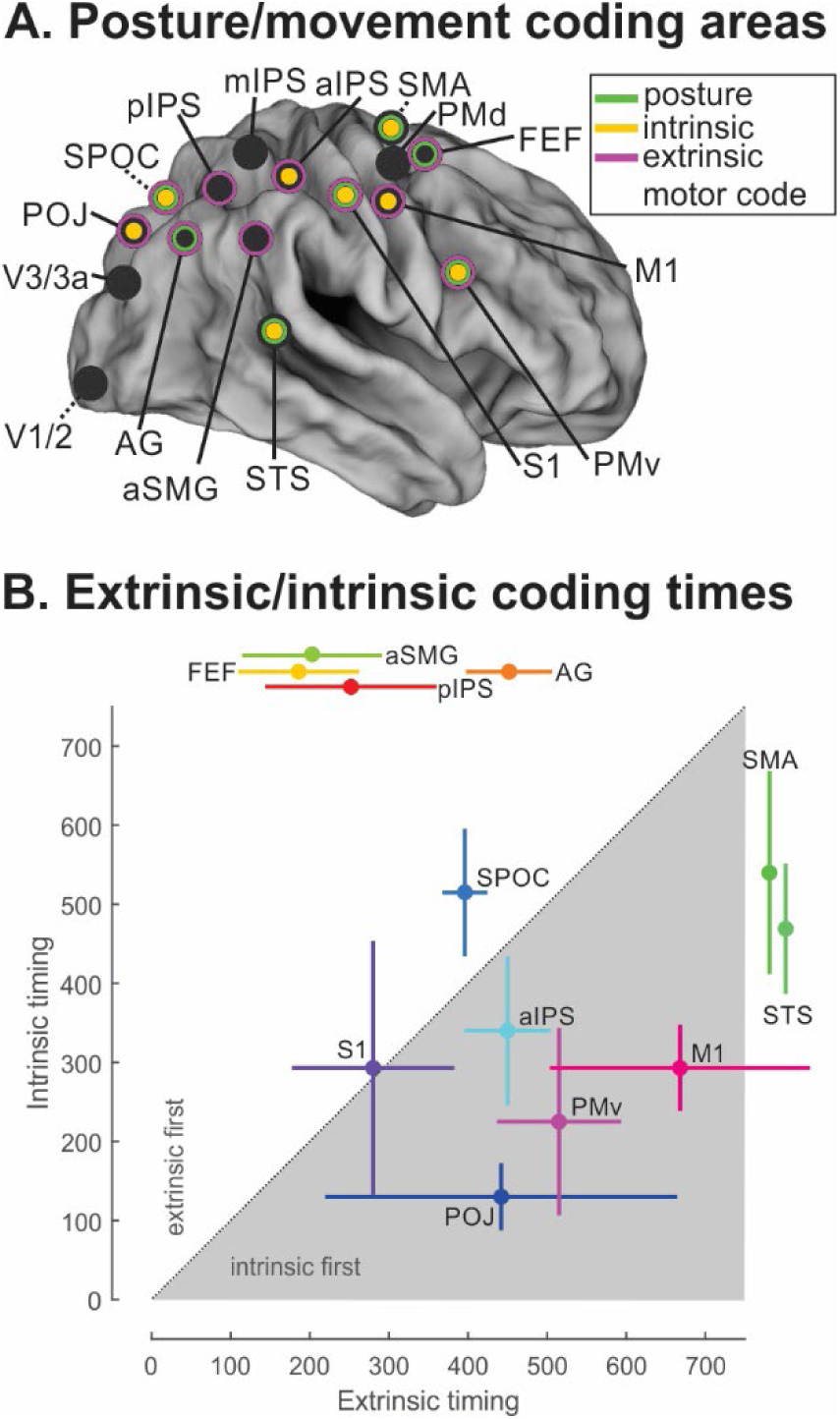
Extrinsic vs intrinsic movement codes. **A**. Summary of coding results across different bilateral ROIs. Colors indicate which areas displayed posture coding, intrinsic and extrinsic motor codes. **B**. Timing of significance (bootstrapped across participants) of intrinsic and extrinsic motor codes between cue onset and (average) movement onset (at 1,815ms after cue onset). Areas showing both intrinsic and extrinsic coding are plotted within the axes (SPOC, M1, S1, POJ, PMv, aIPS). Areas only showing extrinsic (FEF, aSMG, pIPS, AG) or intrinsic (SMA, STS) codes are on the top (horizontal) or right (vertical), respectively. Below diagonal values indicate intrinsic coding first; above diagonal values indicate earlier extrinsic coding. Error bars are 95% confidence intervals.

To better understand that processing sequence of motor planning, we analyzed the relative timing across sites of earliest appearance of significant intrinsic movement and / or extrinsic motor coding. Following the standard logic, one would expect extrinsic coding to appear first, followed by intrinsic coding. Since many brain areas showed both significant extrinsic and intrinsic coding, we plotted the earliest significance times against one another for each brain area. Areas that only showed intrinsic or extrinsic coding, are represented outside the plot as vertical and horizontal lines on the right or top, respectively. The earliest intrinsic coding appeared in POJ at 130ms (±39ms 95% CI); the earliest extrinsic code in FEF at 164ms (±73). Further, of the six areas that coded both intrinsic and extrinsic movement plans, four (POJ, M1, PMv, aIPS) showed earlier intrinsic than extrinsic codes (gray below diagonal area); one showed earlier extrinsic than intrinsic code (SPOC) and one showed simultaneous extrinsic and intrinsic coding (S1). These findings appear to suggest that, contrary to the usual assumption, intrinsic codes can appear *before* extrinsic codes.

## Discussion

We asked when and where in the sensory-motor network posture was coded and integrated into a movement plan to form the required intrinsic motor code. We found posture-related oscillations and extrinsic codes in various frequency bands (from alpha to high gamma), whereas intrinsic codes were specific to alpha and beta bands. Further, we found partially overlapping networks for extrinsic codes (POJ, pIPS, SPOC, AG, aIPS, pIPS, AG, S1, M1, pIPS, SMG, S1, PMv, M1, FEF, aIPS, pIPS) versus intrinsic motor coding (SPOC, POJ, STS, PMv, aIPS; SPOC, SMA, S1, M1). Perhaps most importantly, intrinsic codes were found to arise first and persist in parallel with extrinsic codes.

### Limitations

We did not correct for multiple comparisons beyond temporal clustering which has been shown to abolish false positives in noisy data (Maris & Oostenveld, 2007). Indeed, as expected we found no significant results for sensory coding. Among the remaining 384 TFRs (posture coding and extrinsic and intrinsic motor code) the number of significant responses we found is much higher than expected by chance (5% false positive rate). We are thus confident that the reported significant results are likely to be real, although the same cannot be said for negative results which should be interpreted with caution.

In our experimental design, posture was identical for the whole block of trials to avoid large arm movement-related EMG artifacts. Hence, we could not test the latency of the effects of changes in posture. Since all data was baseline corrected, areas that coded posture independently of movement (such as S1) did therefore not show a posture effect. Thus, the presence of significant posture effects despite baseline correction indicates active involvement of these signals in some specific computation.

Our analysis logic assumes anti-symmetry between Upright and Downward posture motor codes if intrinsic motor plans are present (Figure 2). However, this does not necessarily have to be the case. For example, in probabilistic population codes (Deneve et al., 2001), or in cosine-tuned population MEG responses. Thus again, negative results should be interpreted with caution.

Finally, MEG source localization inherently suffers from limitations in spatial resolution. The better the signal-to-noise (SNR) ratio, the more precise the source localization, but the converse is also true. We have generally found excellent agreement between our source localization results (Blohm et al., 2019) and previous fMRI results (Cappadocia et al., 2017, 2018; Fujiwara et al., 2017), but the precision of our results should still be taken with some caution.

### Networks for posture, extrinsic and intrinsic movement coding

We found that a subset of the visuomotor areas that we previously identified (Blohm et al. 2019) showed posture modulations, extrinsic motor codes, and/or intrinsic motor coding. Our findings of posture modulation and intrinsic coding in areas PMv and M1 during the delay period are generally consistent with the non-human primate literature (Kakei et al., 1999, 2001, 2003; Morrow et al., 2007; Yanai et al., 2008), but the latter point toward mixed coordinates along a posture gain-modulated continuum between extrinsic and intrinsic reference frames. Gain modulation and posture coding could produce the same results in our MEG data, so our study cannot distinguish between these. Our intrinsic coding computations are specific to detecting the spatial (a-)symmetries between left / right extrinsic space vs left / right intrinsic muscle activations. Thus, significant intrinsic coding indicates that there is at least a partial shift of population reference frames from extrinsic to intrinsic codes.

Our previous study (Blohm et al., 2019) averaged MEG data across 3 postures (Upright, Down and Pronation), to identify cortical sites with visuomotor coding. Here, we separated these into partially overlapping networks with extrinsic motor coding (PMv, M1, aIPS, aSMG, POJ, pIPS, SPOC, FEF and AG) versus intrinsic coding (SMA, M1, S1, aIPS, SPOC, POJ, STS and PMv). We did find motor coding in PMd in our previous study, but not in the current study. Otherwise, our previous and current results are generally consistent in the observed location and timing (see below).

Our results are also generally consistent with those of Fujiwara et al., (2017), who used multi-voxel pattern analysis of fMRI data to decode population movement tuning as a function of posture. Like us, they found intrinsic coding in M1 and S1 and no intrinsic coding in PMd and V1. However, they did not find intrinsic coding in PMv. Furthermore, they found intermediate intrinsic-extrinsic coordinates in an area they called ‘PPC’. This area was quite large and could include our areas mIPS, pIPS, and aIPS, which showed different codes. Fujiwara et al., (2017) had inconclusive results in SMA and SOG (superior occipital gyrus, potentially our area V3/V3a), whereas we found a significant intrinsic motor code in SMA and not V3/V3a. Finally, they did not test STS and POJ, which showed intrinsic coding in our study.

### Frequency specificity

MEG allows one to analyze the frequency specificity of task-related signal changes (D. O. Cheyne, 2013). Inhibition shapes the strength of oscillations in the brain (Buzsáki & Watson, 2012; Sherfey et al., 2018), and inhibition in long-range cortical-thalamic loops, medium-range cortico-cortical coupling and short-range neuron-to-neuron interactions have been associated with alpha (Suffczynski et al., 2001), beta (Cabral et al., 2014) and gamma (Fernandez-Ruiz et al., 2023; Jadi & Sejnowski, 2014) oscillations, respectively (Buzsáki & Wang, 2012; Ray et al., 2008). These frequency bands have been associated with different sensory-motor and cognitive processes. Changes in alpha power reveal sensory processing (Buchholz et al., 2014; Jensen et al., 2002; Klimesch, 2012; Palva & Palva, 2007); beta power correlates with aspects of sensory-motor planning and control (Buchholz et al., 2014; D. O. Cheyne, 2013; Isabella et al., 2015; Kilavik et al., 2013; Lopes da Silva, 2013; Neuper et al., 2006; Neuper & Pfurtscheller, 2001; Spitzer & Haegens, 2017; Van Der Werf et al., 2009); and gamma power reflects memory activity across different putative brain functions (Buzsáki & Wang, 2012; Düzel et al., 2010; Ray et al., 2008).

We found changes in oscillatory power across all frequency bands suggesting that posture and extrinsic coding were processed (alpha) actively maintained (gamma) and used in the sensory-to-motor transformation to create an intrinsic movement plan (beta). Surprisingly, intrinsic movement coding was present not just in the beta band but also in the alpha band, even at later times during the delay period. We speculate that this reflects long-ranging network interactions in establishing an intrinsic motor plan.

POJ and SPOC showed both extrinsic and intrinsic coding in the same band, i.e. alpha and beta band, respectively. For SPOC, while there was overlap in motor coding schemes in beta band, extrinsic coding was observed earlier and then seemed to switch to intrinsic coding later, prior to the movement onset. POJ had a period of earlier joint extrinsic and intrinsic coding, but then only sustained the intrinsic code for longer. This contrasted with S1, M1, PMv and aIPS who all coded extrinsic and intrinsic information simultaneously but in different frequency bands.

### Timing

We previously reported (Blohm et al., 2019) that planning activity is first dominated by a bottom-up sweep of sensory activity from occipital to parietal and frontal areas, followed by a top-down progression of motor coding from frontal to posterior areas. Only much later during the delay period did we then find hand-specific motor coding (Blohm et al., 2022). Our current results are somewhat at odds with this interpretation.

Here — by disentangling intrinsic and extrinsic coding — we found that both could be decoded as early as ∼180ms after cue onset, much earlier than previously reported. Evidence for intrinsic motor plans was first found in areas POJ and S1, then gradually in other areas like PMv, STS, aIPS, SMA, and SPOC, and finally M1. Extrinsic motor plans first appeared in FEF and then in S1, POJ, SPOC, M1, aSMG, aIPS, AG, pIPS and PMv. In areas showing both evidence for extrinsic and intrinsic coding, intrinsic was first in POJ, S1, aIPS and PMv; and extrinsic coding was first in SPOC and M1. Overall, extrinsic coding appeared ∼30ms after intrinsic coding (∼160ms if only considering areas coding for both). Further, if we removed FEF (which could be reflecting oculomotor signals) intrinsic motor coding would average 50ms earlier than extrinsic motor coding. This suggests that the sensorimotor transformation first establishes an early intrinsic motor plan in parietal and frontal planning areas and then computes a more general extrinsic movement plan.

Our finding that intrinsic movement plans are established before (or at least in parallel to) extrinsic codes means that one cannot think about sensorimotor transformations as a serial process (Blohm et al., 2009). Instead, all relevant signals might be used for the primary goal of sensorimotor transformation: rapid conversion of sensory inputs into motor plans appropriate for muscle activation. This suggests that muscle space may be relevant in planning areas. Conversely, extrinsic coordinates may be more important later for perception, spatial cognition, predictive consequences of potential future actions, or updating body state. To summarize, our findings suggest a fundamental shift in the way one needs to consider the location and timing of intrinsic and extrinsic movement plans.

## Acknowledgements

Experiments were supported by a Canadian Institutes for Health Research Grant held by JDC. GB was supported by a Marie Curie Fellowship (EU) during the experiments and by NSERC (Canada) thereafter. During these studies, JDC was supported by a Canada Research Chair. The authors would like to thank Andreea Bostan, William Gaetz, Herbert Goltz and Sonja Bells for technical assistance during data collection.

## Footnotes

1 By “coding” we simply mean that there is some signal or modulation of activity that allows us to decode information. We do not imply that there is a specific code or explicit representation of this information.

2 mIPS showed mixed extrinsic / intrinsic coding (Fujiwara et al., 2017)

